# Antifungal activity of *Conocephalum conicum*(L) Dumort. (Marchantiophyta) against *Fusarium oxysporum* f. sp. *lycopersici*

**DOI:** 10.1101/2021.07.27.454003

**Authors:** Kavita Negi, Preeti Chaturvedi

**Affiliations:** Drug Standardization Research Unit, Central Council for Research in Unani Medicine, Ministry of AYUSH, New Delhi; Department of Biological Sciences, College of Basic Sciences and Humanities, G.B. Pant University of Agriculture & Technology, Pantnagar, Uttarakhand, India

**Keywords:** Bryophytes, Biocontrol, *Conocephalum conicum*, *Fusarium oxysporum* f. sp*lycopersici*, Antifungal, SEM and GC-MS

## Abstract

Tomato, a high valuevegetable crop, suffers huge production losses in tropics due to a wilt disease caused by *Fusarium oxysporum*f. sp.*lycopersici*. Present study was undertaken to find an effective biocontrol method to check fusarium wilt in order to curb the losses suffered by the crop growers. Organic extracts(acetone, methanol/ethanol) of thalloid bryophytes (*Conocephalumconicum* (L.) Dumort. and*Marchantiapapillata*Raddi subsp. *grossibarba*(Steph.) Bischl.)were tested against *F. oxysporum* f. sp. *lycopersici*using disc diffusion and micro broth dilution assay.Methanol extract of *C.conicum* (L.) Dumort. (CCDM) showed significantly high antifungal activity (85.5% mycelial inhibition; 31.25μg/mL MIC and 125μg/mL MFC).Potential of methanol extract was tested in a glasshouse experiment on tomato, which illustrated the efficacy of the plant extract to control the fusarial wilt. Morphological and ultrastructural alterationsin CCDM treated fusarium myceliawere observed in scanning electron microscopy. GC-MS analysis of CCDM extract showed the presence of51 constituents, and the dominant compounds werebis (bibenzyl), acyclic alkanes, fatty acids, sesquiterpenpoids and steroids. The study suggested that *C. conicum* being an efficient source of Riccardin C like antifungal compounds provides a potent and eco-friendly alternative to conventional fungicides in vegetables.

## 1. Introduction

Tomato(*Solanum lycopersicum* L.)is used around the world due to its flavour, colour, anti-oxidative and anticancer properties**(Gerszberg and Konka2017)**.It is the world’s largest consumed vegetable crop after potato, sweet potato, and onion and, it also tops the list of canned vegetables **(Olaniyi et al. 2010)**. Presence of lycopene in tomato gives specific benefits against number of ailments.

Fusarium wilt, one of the mostprevalent and widespread diseases of tomato, severely damagesthe crop as *Fusariumoxysporum* f. sp*lycopersici*(FOL)startsgrowingwithin the vascular tissues of the plantand impedes water supply to different parts of the plant**((Ignjatov et al. 2012**, **Moretti et al. 2008)**. **Singh and Kamal (2012)** reported 10-90% loss in tomato yield due to this wilt. Significant losses in tomatoproduction in greenhouse condition and in fields have been reported by **Amini and Sidovich (2010)**.

Carbendazime and propiconazole are some common fungicides which are highly effective against fusarium wilt **(Manasa et al. 2017)**. Fungicides are synthetic in nature and may affect environment and human health adversely due to their toxicity, carcinogenicity and persistent nature **(Fisher et al. 2018)**. Therefore, biocontrol measures seem to be a suitable and sustainable solution to control fusarium wilt. Using botanicals in controlling fusarium wilt provides an ecofriendly, cost-effective and non-toxic biocontrol method **(Trda et al. 2019)**. Botanicals belonging to different plant groups*viz*., lichens and angiospermshave been reported to possess antifungal efficacy against *F. oxysporum* **(Basile et al. 2015; Yeole et al. 2016)**.Antifungal activities have also been reported in some bryophytes(**Tadesse 2002; Kirisanth et al. 2020; Commisso et al. 2021)**.

Bryophyta, a very primitive and simplest group of the embryophytes, have acquiredunique survival strategies. They lack protection shields like bark but possess variety of bioactive compounds against inhospitable environments. Rich repository of these secondary metabolites *viz.,*benzyls, bis benzyl derivatives, sesquiterpenes, phenols etc. in bryophytes have protected them against microbes and, hence, enabled their survival in diverse habitats **(Asakawa, 2017)**. In view of the above, an attempt was made in the present study to assess the efficacy of thalloidbryophytes for an effective control of *F. oxysporum* f. sp*lycopersici*employing *in vitro* and *in vivo* approaches.

## 2. Material and Methods

### 2.1 Collection of plant material

Based on the ethnomedicinal value reported by **Glime (2017)** and **Negi et al(2018)**, two species of bryophytes (liverworts),*Conocephalumconicum* (L.) Dumort(Conocephalaceae) and *Marchantiapapillata*Raddi subsp. *grossibarba*(Steph.) Bischl.(Marchantiaceae) were collected from an altitudinal range of 200-2100m from Uttarakhand, India.*C. conicum* was collected from Mukteshwar (Altitude: 2100m, shady wall) and Dwarahat (Altitude: 1400m, moist wall). *M. papillata*was collected from Dwarahat (Altitude: 1400m, river basin) and Pantnagar (Altitude: 213m, moist wall).Voucher specimens of the collected species (NC501, NC502) were submitted to G.B. Pant University Herbarium, Pantnagar.

### 2.2 Preparation of plant extract

Collected plant samples were thoroughly washed under running tap water, shade dried, pulverized and extracted in 80 % ethanol/methanol and acetone (Analytical Grade, Sigma Aldrich). Extraction was performed using hot (Soxhlet) and cold percolation methods (10 g/100 mL). The extracts were filtered through Whatman No. 1 filter paper and concentrated using a rotary evaporator(Biogen Scientific). The crude extract was dissolved in the respective solventfor the preparation of stock solution of 1mg/mL. The dilutions of different concentrations (100, 400, 700, 1000 μg/mL) were prepared from the stock solution and used for further study.

### 2.3 Antifungal Assay

*Fusarium oxysporum*f. sp. *lycopersici*(FOL) was obtained from the Rhizosphere laboratory of Department of Biological Sciences, G.B. Pant University of Agriculture & Technology, Pantnagar**(Joshi et al. 2013)**. The culture was revived on Potato dextrose agar(Himedia) at 27 °C and the same was used for disc diffusion assay. For further experiments, inoculum was prepared by culturing FOL in potato dextrose broth in shake culture at same temperature for 5-6 days.

For disc diffusion assay, Potato Dextrose Agar (PDA) medium was poured aseptically in the Petri plates (90 mm) and plates were allowed to solidify.Extract (20 μL) of varying concentrations in different solvents was pipetted out into the discs (Hi Media). Four discs (5 mm), two treated (T) with plant extract (20μl) and two control (C) (20μl) along with the test fungus(as shown in **Fig 1)** were kept in the same Petri plate following a standard protocol of **Bauer (1966)** with minor modifications as suggested by **Negi and Chaturvedi (2016a)**.Carbendazim (Bavistin 50 %) was used as a positive control while the solvents were used as negative control in different plates. Inoculated fungal plates were incubated for 5 days at 28± 2° C. Inhibition (%) of fungal growth was calculated.

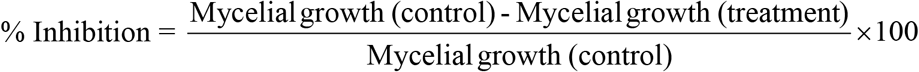

**FIGURE 1.** Plate ofdisc diffusion Assay. C = Control disc; T = Treatment disc; F=*Fusarium oxysporum*

### 2.4 Determination of Minimum Inhibitory/Fungicidal concentration

Micro broth dilution assay was performed to determine the inhibitory and fungicidal concentration of bryophyte extracts using freshly prepared potato dextrose broth (PDB) as diluents **(Janovska et al. 2003)**. Fresh and revived culture of the test fungus was diluted upto100 folds in broth (100μl of microorganism in 10 mL broth). Colony forming units (CFU) of the fungus were determined (1×10^9^ CFU/mL) by taking OD at 620 nm using a UV-Visible spectrophotometer (Thermo Fisher Scientific)**(Sutton 2011)**. Plant extracts, with 1000 to 0.98 μg/mL concentration in two-fold dilution series were added to the test tubes containing fresh fungal cultures. All the tubes with fungal cultures were incubated at 28°C for 72 h. The visible turbidity and optical density of the cultures were determined at 620 nm using spectrophotometer. The lowest concentration of the extract that inhibited the visible growth of the test microorganisms was recorded as minimum inhibitory concentration (MIC) and the test culture without any visible microbial growth was considered as minimum fungicidal concentration (MFC).

### 2.5 Soil collection and preparation of experimental pots

Autoclaved loamy soil and river bed sand were used (sand: soil= 3:1)in the pot experiment. The pH and EC of this mixture were 8.15 and 65.7 μS, respectively. Tomato seeds were surface sterilized in 1% HgCl_2_ for 2-3 min, rinsed three times in sterile distilled water and then dipped in sterile water for 1 day for imbibition. After drying in towel paper, the seeds were sown in the sterilized soil mixture in the Departmental glasshouse. Twenty days old tomato seedlings (four leaf stage) grown in autoclaved sand: soil (3:1)were transplanted into 500g pot containing similar ratio of sand and soil. Plants were irrigated with tap water and left for 1-2 days for equilibration before setting up the experiment. Thefungalinoculum was taken from PDB as it contained a large number of macroconidia. The population of FOLin the substrate was estimated asthenumber of CFU per gram of soil. Approximately 2 ×10^8^ CFU/g of the fungal inoculum was used for pathogenesis in tomato roots.1 ml of fungal inoculums suspension (made in water) was used for inoculating the seedlings by soil drenching method.Theinfection of FOL was confirmed by the symptoms of wilting in tomato plants within 15 days of the experiment.

Pot experiment was conducted in a completely randomized block designin the glass housewith 5 replicates each using two different treatments*viz*., 125 μg/mL and 31.25 μg/mL of aqueous solution of CCDM extract (EC1, EC2 respectively). All the treatments were applied in form of fine solution to the roots of tomato plants per pot until the roots absorbed the solution completely. The treatments were given one day prior and 15 days after FOL treatment. Three control(s) used were fusarium infested negative control (C+F), non-infested water control, and positive control (carbendazim). Pots were irrigated with deionised water as and whenrequired.

Pots were maintained in the glasshouse under the following growth conditions: Temperature −27°C, Photoperiod-16/8 hour day/night cycle, Light intensity-400 Em-2s-^1^, (400-700 nm), Relative humidity-60%.

### 2.6 Observations

Observations related to shoot length, root length, fresh weight and dry weight of tomato plants were taken for a period of 30 days after planting.After harvesting the plants, roots were washed thoroughly with tap water followed by washing with deionised water. Shoot and root length were measured from the soil base to the tip of the fully expanded leaf and soil base to the tip of the root. Shoot and root fresh weight (g) were weighed separately immediately after harvesting. Dry weight of shoot and roots were taken (g) separately after drying the samples at 60°C inan electric oven for 48 h.

### 2.7 Gas Chromatography-Mass Spectroscopy (GC-MS)

The crude methanol extract of *C. conicum*(CCDM)was filtered using 0.45μm syringe filters. GC-MS analysis of the extract was carried out using a GC-MS System (Shimadzu) QP 2010 equipped, with Rxi ^®^-5Sil MS capillary GC column (5 % phenyl 95 % dimethyl polysiloxane) with 30 m length, 0.25 mm diameter and 0.25 μm film thickness at Central Instrumentation Facility, Jawaharlal Nehru University, New Delhi. Helium was the carrier gas and used at a flow rate of 16.3 mL/min. Sampling time was maintained at 1 min.at a column pressure of 81.7 kPa. Column oven and injector temperatures were maintained at 80 ° C and 270 ° C respectively. Samples (1μl) were injected into the column with a split ratio of 10:1. Names, molecular weight and structures of the individual compounds were identified by matching their mass fragmentation pattern with the National Institute of Standard Technology (NIST) Library.

### 2.8 Scanning electron microscopic (SEM) study

Effect of methanol extract of *C. conicum* on FOLwas observed by SEM following a standard protocol of **Plodpai et al. (2013)** with minor modifications as reported by **Negi and Chaturvedi (2016a)**. SEM (JEOL6610LV) facility was provided by Department of Anatomy, College of Veterinary and Animal Sciences, G.B. Pant University of Agriculture & Technology, Pantnagar, India.

### 2.9 Statistical Analysis

Analysis of variance was calculated using standard statistical methods **(Snedecor and Cochran 1994)**. Disc diffusion assay was performed in triplicate while pot experiment was conducted using five replicates. Values were represented as mean ±SEm.Mean value comparison was computed usingfour factorial ANOVA (for antifungal activity through disc diffusion assay). It revealed level of significance at *P*<0.05 and *P*<0.01 among different bryophyte species, solvents, concentrations and days. The analysis was carried out using IBM SPSS statistical software, version 15.0 (IBM, New York, USA).

## 3. Results

### 3.1 Bioactivity of plant extract against *Fusariumoxysporum*f. sp. *lycopersici*

Organic extracts (alcohol and acetone) of *Conocephalum conicum*and *Marchantia papillata* showed significant antifusarium activity (Fig 2). Organic extracts (100-1000 μg/mL) of bryophytes revealed dose - dependent inhibition of FOL (using disc diffusion assay) at different time intervals (2-5 days) (Table 1 and Fig 2). The methanol extract of *C. conicum* (CCDM) showed maximum per cent inhibition of FOL mycelial growth, values ranging from 25.31±0.005 (after 2 days) to 85.5±0.57 (after 5 days) at the concentration of 100 and 1000 μg/mL respectively (Table 1). Similarly, acetone extract of *C. conicum* (CCDA) showed second highest per cent inhibition from 31.4±0.57(after 2 days) to 70.16±0.5 (after 5 days) at 100 and 1000μg/mLrespectively. Minimum inhibitory concentration (MIC) and minimum fungicidal concentration (MFC) of different organic extracts of bryophytes ranged from 31.25 to 500μg/mL and 125 to 500 μg/mL respectively (Table 2). Again, CCDM extract was found most potent against FOL with lowest MIC (31.25μg/mL) and MFC (125 μg/mL). Hence, the effectiveness of CCDM extract was further assessed by chemical characterization and *in vivo* pot experiment.

**FIGURE 2.**
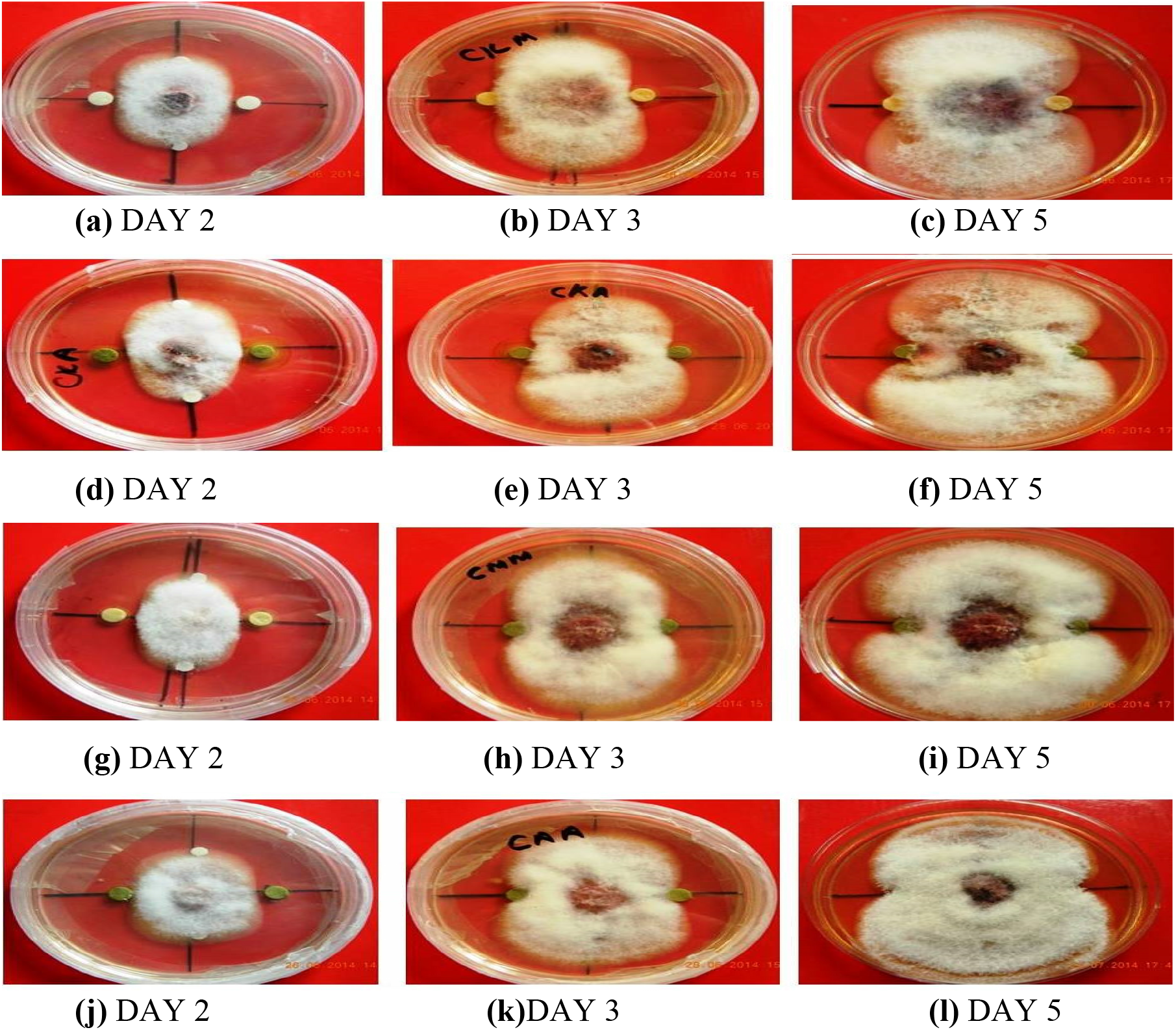
Disc diffusion plates showing growth inhibition of *Fusarium oxysporum*f. sp*. lycopersici* under different treatments. **(a, b, c)** Mycelia treated with CCDM extract, **(d, e, f)** Mycelia treated with CCDA extract,**(g, h, i)** Mycelia treated with CCMM extract,**(j, k, l)**Mycelia treated with CCMA extract (CCDM-methanol extract of *Conocephalum conicum* collected from Dwarahat, CCDA-acetone extract of *C. conicum* from Dwarahat, CCMM-methanol extract of *C. conicum* from Mukteshwar, CCMA-acetone extract of *C. conicum* from Mukteshwar).

**Table 1.** Growth inhibition (%) of *Fusarium oxysporum*f. sp*. lycopersici*(FOL)with different extracts of bryophytes

| Name of<br>Bryophyte<br>extracts | Percent Inhibition (%) |  |  |  |  |  |  |  |  |  |  |  |  |  |  |
| --- | --- | --- | --- | --- | --- | --- | --- | --- | --- | --- | --- | --- | --- | --- | --- |
|  | Conc. (100µg/mL) |  |  | Conc. (400µg/mL) |  |  | Conc. (700 µg/mL) |  |  | Conc. (1000 µg/mL) |  |  |  |  |  |
|  | Day 2 | Day 3 | Day 5 | Day 2 | Day 3 | Day 5 | Day 2 | Day 3 | Day 5 | Day 2 | Day 3 | Day 5 |  |  |  |
| CCDM | 25.31±0.05 | 26.86±0.05 | 47.84±0.57 | 50.84±0.57 | 44.13±0.56 | 51.35±0.55 | 66.10±0.28 | 67.58±1.14 | 75.12±1.1 | 82.03±0.5 | 84.4±1.1 | 85.5±0.57 |  |  |  |
| CCDA | 31.4±0.57 | 49.34±0.57 | 42.86±0.57 | 49.58±0.57 | 41.09±0.57 | 49.72±1.15 | 60.95±0.57 | 66.42±0.66 | 64.91±0.55 | 65.28±0.57 | 66.43±0.57 | 70.16±0.57 |  |  |  |
| CCMM | 22.43±0.88 | 27.23±0.57 | 27.45±0.57 | 53.84±0.57 | 61.63±0.33 | 43.88±0.57 | 58.08±1.1 | 59.56±0.28 | 61.36±0.57 | 56.25±0.57 | 62.08±1.11 | 62.79±0.57 |  |  |  |
| CCMA | 23.63±0.33 | 25.54±0.57 | 24.89±0.57 | 32.5±0.57 | 30.95±0.66 | 41.07±0.33 | 17.18±0.33 | 43.43±0.57 | 49.10±0.66 | 47.5±0.57 | 35.71±0.57 | 58.23±0.57 |  |  |  |
| MPDE | 20.6±0.57 | 23.33±0.57 | 21.54±0.57 | 46.80±0.57 | 50.88±0.88 | 49.15±0.66 | 36.09±0.66 | 51.6±0.33 | 49.74±0.57 | 31.66±0.57 | 45.61±0.57 | 55.97±0.33 |  |  |  |
| MPDA | 12.56±0.66 | 14.71±0.57 | 14.34±0.57 | 12.54±0.57 | 13.63±0.28 | 34.73±0.57 | 29.26±0.33 | 32.63±0.33 | 41.02±0.33 | 46.58±0.33 | 49.22±0.57 | 54.02±0.33 |  |  |  |
| MPPE | 12.48±1.1 | 14.48±0.66 | 14.48±1.15 | 18.64±0.33 | 56.25±0.33 | 43.95±0.33 | 24±0.66 | 40.47±0.66 | 44.96±0.57 | 42.27±1.15 | 40.88±0.66 | 53.97±0.33 |  |  |  |
| MPPA | 11.23±0.33 | 13.56±0.66 | 14.67±1.15 | 16.45±0.66 | 51±0.88 | 41.89±0.33 | 20±0.66 | 36.89±0.66 | 41.96±0.5 | 40.27±1.1 | 39±0.66 | 51.89±0.33 |  |  |  |
| Carbendazim | 72.0±0.66 | 78.48±0.33 | 88.87±1.1 | ND | ND | ND | ND | ND | ND | ND | ND | ND |  |  |  |
|  | Bryophyte species | Organic solvent | Conc | Days |  |  |  |  |  |  |  |  |  |  |  |
|  | (A) | (B) | (C) | (D) | AB | AC | AD | BC | BD | CD | ABC | ABD | ACD | BCD | ABCD |
| CD at 1% | 0.39 | 0.27 | 0.39 | 0.34 | 0.55 | 0.79 | 0.68 | 0.96 | 0.48 | 0.68 | 1.11 | 0.96 | 1.37 | 0.96 | 1.9 |
| CD at 5% | 0.30 | 0.21 | 0.30 | 0.26 | 0.42 | 0.60 | 0.52 | 0.73 | 0.36 | 0.52 | 0.85 | 0.73 | 1.04 | 0.73 | 1.47 |
| SEm | 0.10 | 0.07 | 0.10 | 0.09 | 0.15 | 0.21 | 0.18 | 0.26 | 0.13 | 0.18 | 0.30 | 0.26 | 0.37 | 0.26 | 0.53 |

*CCDM-methanol extract of Conocephalum conicum from Dwarahat, CCDA- acetone extract of C. conicum from Dwarahat, CCMM-methanol extract of C. conicum from Mukteshwar, CCMA-acetone extract of C. conicum from Mukteshwar, MPDE- ethanol extract of Marchantiapapillata from Dwarahat, MPDA- acetone extract of M. papillata from Dwarahat, MPPE-ethanol extract of M. papillata from Pantnagar, MPPA-acetone extract of M. papillata from Pantnagar, Conc.-Concentration, ND-not determined. All the experiments were performed in triplicates. Mean value (±SEm) with four factorial ANOVA revealed level of significance at $P<0.05$ and $P<0.01$ among different bryophyte species, solvents, concentrations and days.*

**Table 2.** Minimum Inhibitory Concentrations (MIC) and Minimum Fungicidal Concentrations (MFC) of different extracts of bryophytes against *Fusarium oxysporum*f. sp.*lycopersici.*

| Name of Bryophyte extracts | Minimum Inhibitory Concentration MIC (µg/mL) | Minimum Fungicidal Concentration MFC (µg/mL) |
| --- | --- | --- |
| CCDM | 31.25 | 125 |
| CCDA | 62.5 | 250 |
| CCMM | 62.5 | 250 |
| CCMA | 125 | 500 |
| MPDE | 125 | 500 |
| MPDA | 125 | - |
| MPPE | 250 | - |
| MPPA | 500 | - |

*CCDM-methanol extract of Conocephalum conicum from Dwarahat, CCDA- acetone extract of C. conicum from Dwarahat, CCMM-methanol extract of C. conicum from Mukteshwar, CCMA- acetone extract of C. conicum from Mukteshwar, MPDE- ethanol extract of Marchantiapapillata from Dwarahat, MPDA- acetone extract of M. papillata from Dwarahat, MPPE-ethanol extract of M. papillata from Pantnagar, MPPA-acetone extract of M. papillata from Pantnagar, (-)=did not show fungicidal concentration.*

### 3.2 Chemical Characterization

The most potent extract (CCDM) was chemically characterized using gas chromatography and mass spectrophotometry. Percentage and the retention timings of the major components are given in Table 3 and Fig 3. It showed the presence of 51 constituents contributing 100% of the total extract. The results revealed that the extract was dominated by bis (bibenzyl) = riccardin c (64%), acyclic alkanes (eicosane = 5%), fatty acids (n-hexadecanoic acid = 3.43%), sesquiterpenpoids (10-epi-α-eudesmol = 1.6%) and steroids (δ.5-ergostenol = 4.3%, stigmasterol = 1.2%).

**Table 3.**
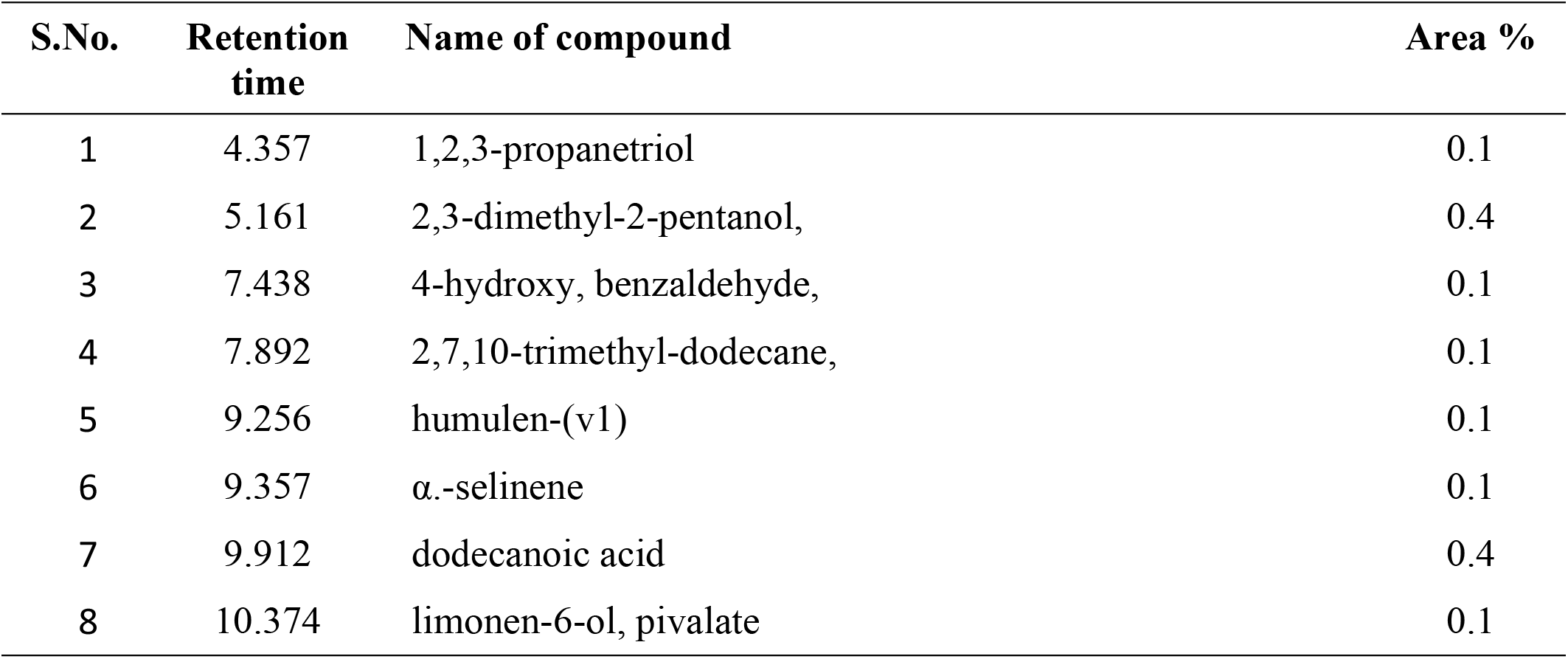
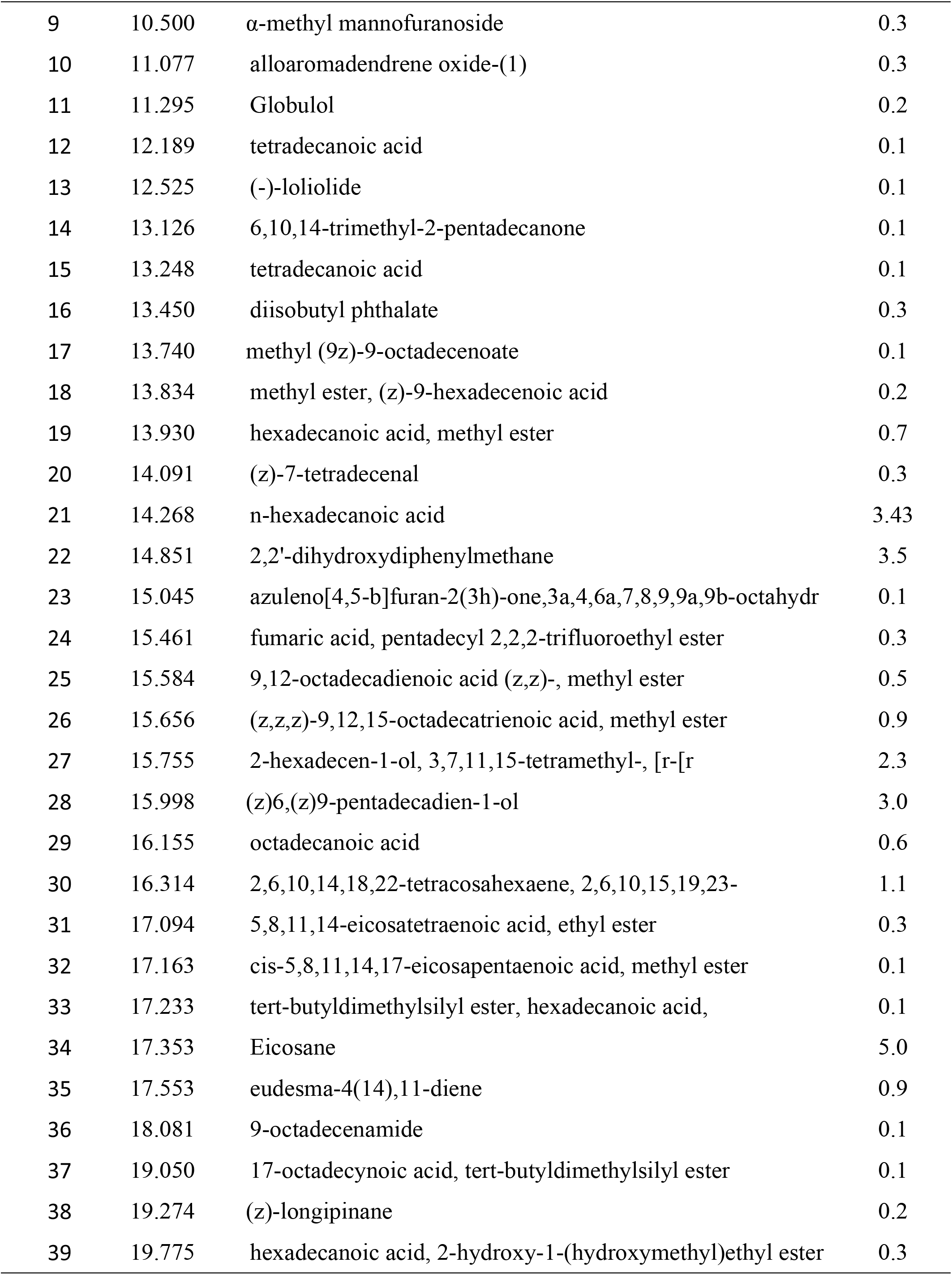
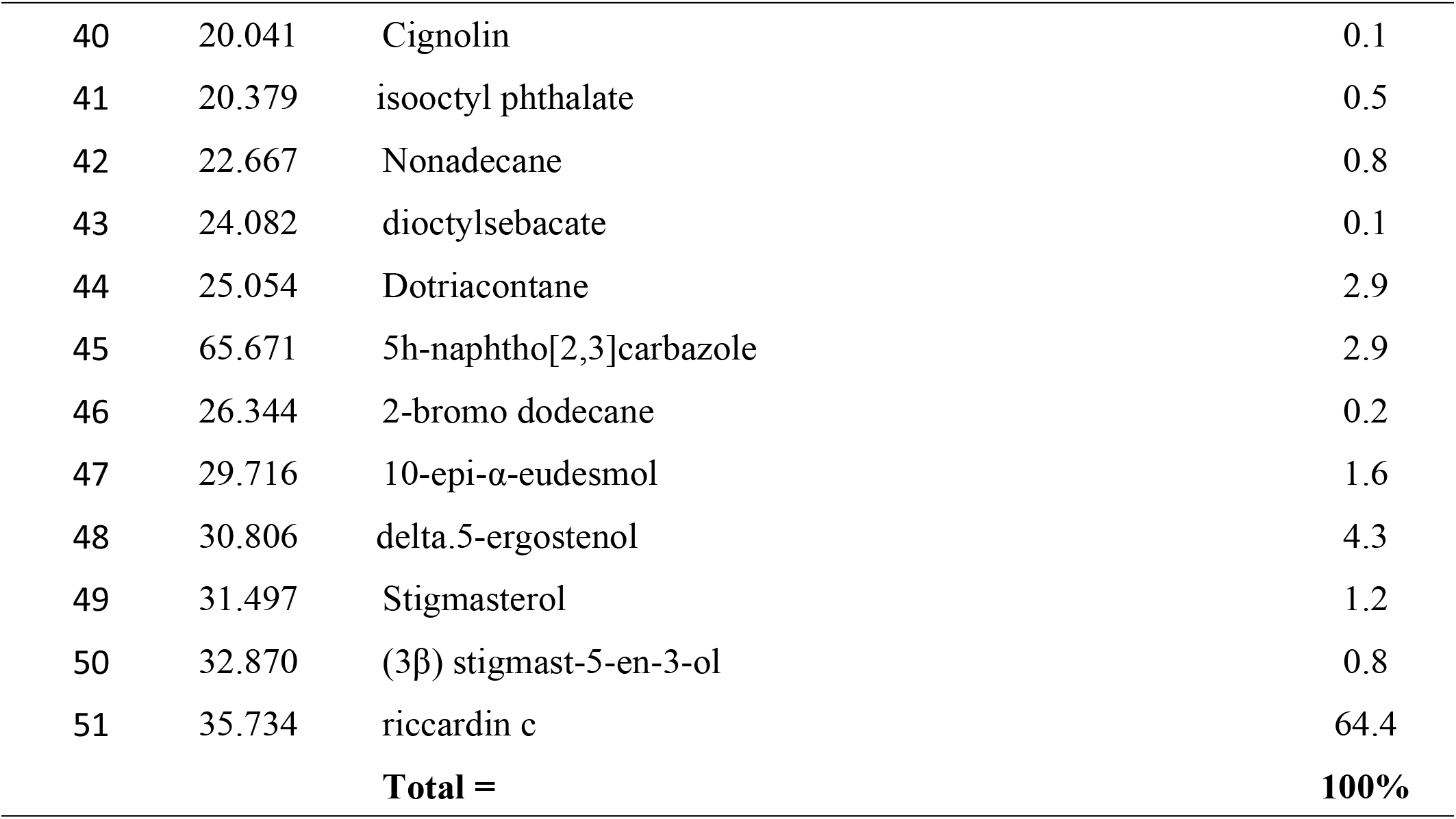
Chemical characterization (Gas Chromatography-Mass Spectrometry) of CCDM extracts (methanol extract of *Conocephalumconicum* collected from Dwarahat).

**FIGURE 3.**
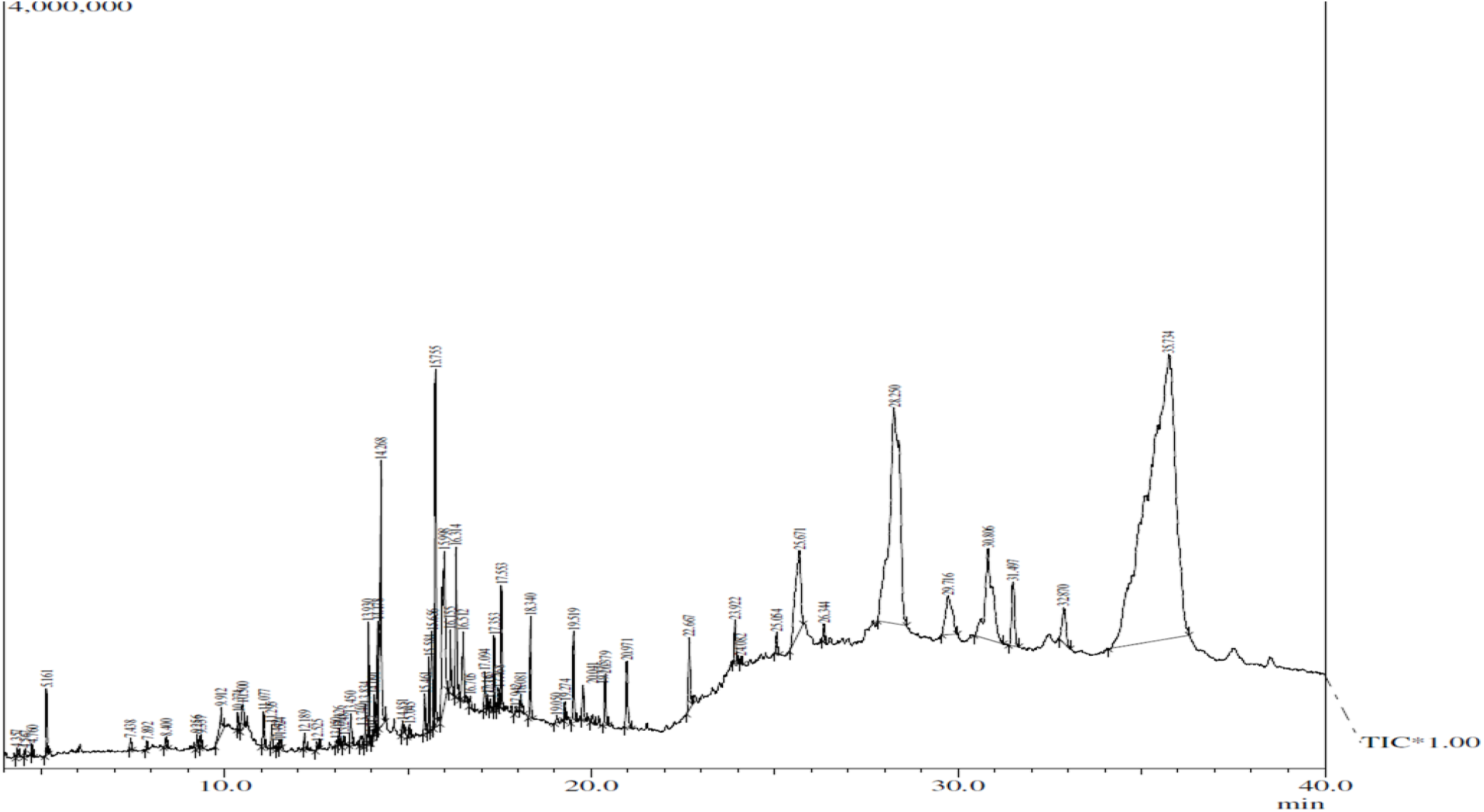
GC–MS chromatogram of methanol extract of *Conocephalum conicum* Dwarahat (CCDM)

### 3.3 Effect of CCDM extract on hyphal and conidial morphology

Scanning electron micrographs (SEM) demonstrated that 125 μg/ml (EC1) of CCDM extract caused profound changes in hyphal morphology of FOL (Fig 4). Treated mycelia appeared wrinkled and ruptured. In contrast, SEM image of untreated fungus mycelia displayed smooth hyphal surface and intact conidia with typically tapered apices.

**FIGURE 4.**
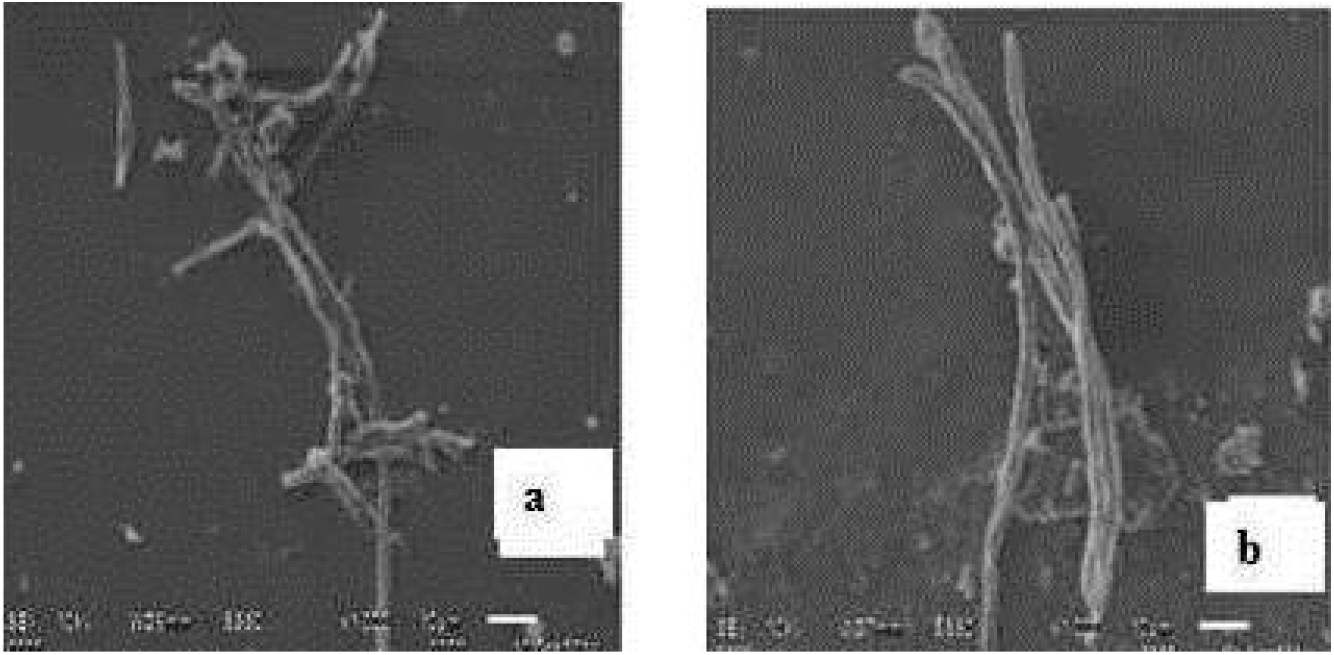
SEM imagesof (a) ruptured mycelia under EC1 treatment of CCDM extract (b) mycelia in control

### 3.4 Effect of CCDM extract on growth parameters of tomato plants

CCDM extract was tested for *in vivo* antifungal efficacy against FOL in a pot experiment using two most effective concentrations, EC1 (125 μg/mL) and EC2 (31.25μg/mL) besides keeping a positive control (carbendazim treated plants + Fusarium inoculated), a negative control (Fusarium inoculated without any treatment) and a water control in the pot experiment (30 days). Results of the experiment are given in Fig 5, Fig 8.

#### 3.4.1 Observations of shoot and root length

Both EC1 and EC2 treated plants showed significantly higher shoot/root length compared to the negative control (Fusarium inoculated without any treatment) after 15 and 30days of the treatment (Fig 5a, 5b). After 30days, the EC1 treated plants showed comparable shoot length (35.50±0.36cm) and root length (15.17±0.27cm) as that of the positive control (36.77±0.22cm; 16.27±0.12cm). Shoot and root length was, however, significantly low in all other inoculated treatments compared to water control.

**FIGURE 5a.**
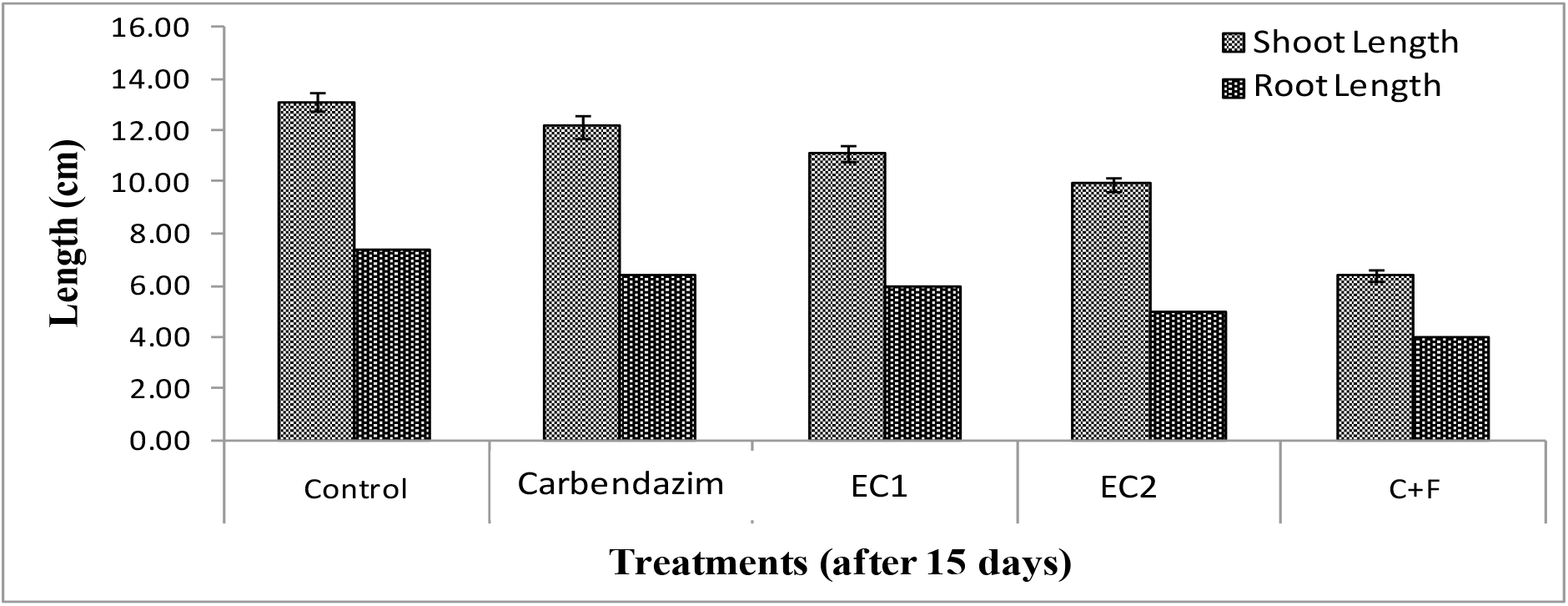
Effect of different doses of CCDM extract on shoot/ root length of tomato plants(after 15 days). Control= water; Carbendazim = positive control; EC1 = 125μg/mL; EC2 = 31.25μg/mL; C+F = negative control (*Fusarium* infested without treatment). Data represents mean ±SE from five replicates

**FIGURE 5b.**
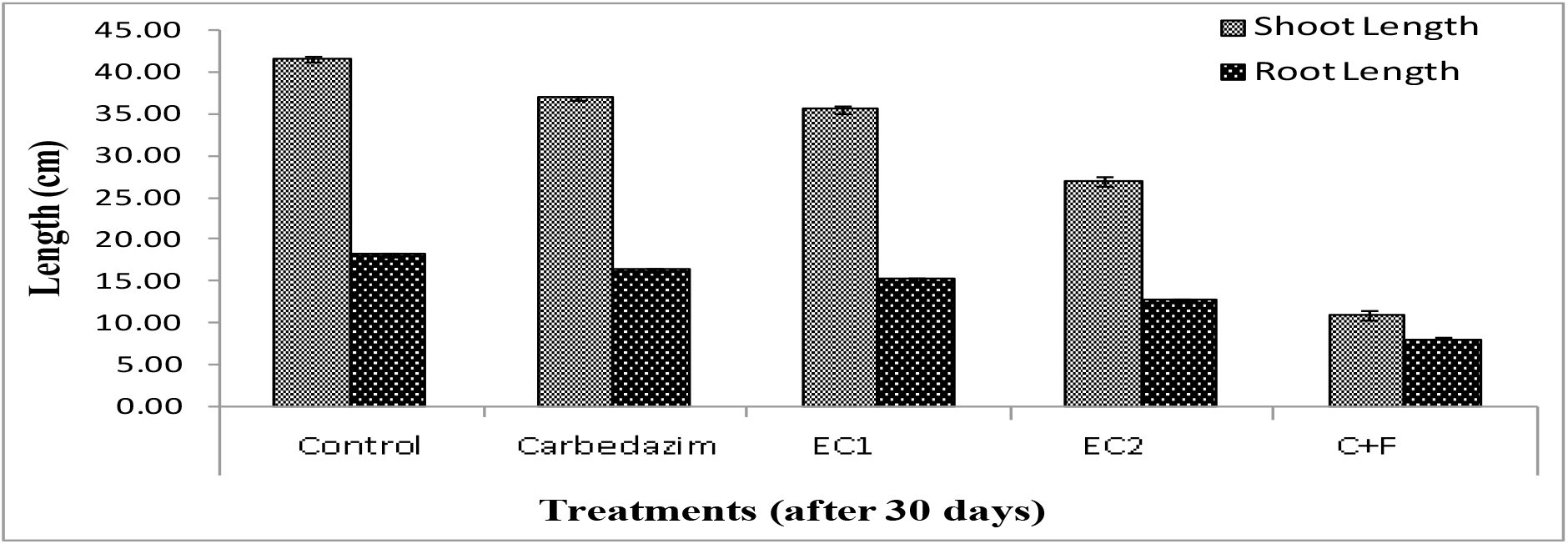
Effect of different doses of CCDM extract on shoot/root length of tomato plants (after 30 days). Control=water, Carbendazim = positive control; EC1 = 125μg/mL; EC2 = 31.25μg/mL; C+F = negative control (*Fusarium* infested plant without treatment). Data represents mean ±SE from five replicates

#### 3.4.2 Shoot and root fresh weight

A similar trend of shoot and root fresh weight (with different treatments) was observed in the pot experiment after 15 and 30 days of setting up the experiment (Fig 6a, Fig 6b). After 30 days, EC1 treated plants showed shoot and root fresh weight in EC1 treated plant was 4.62±0.18 g and 2.26±0.03grespectively after 30 days, where as inpositive control it was 4.91±0.20g and 2.67±0.01g after same time. Reduction of wilt in EC2 treated plants was significantly higher than the negative control.

**FIGURE 6a.**
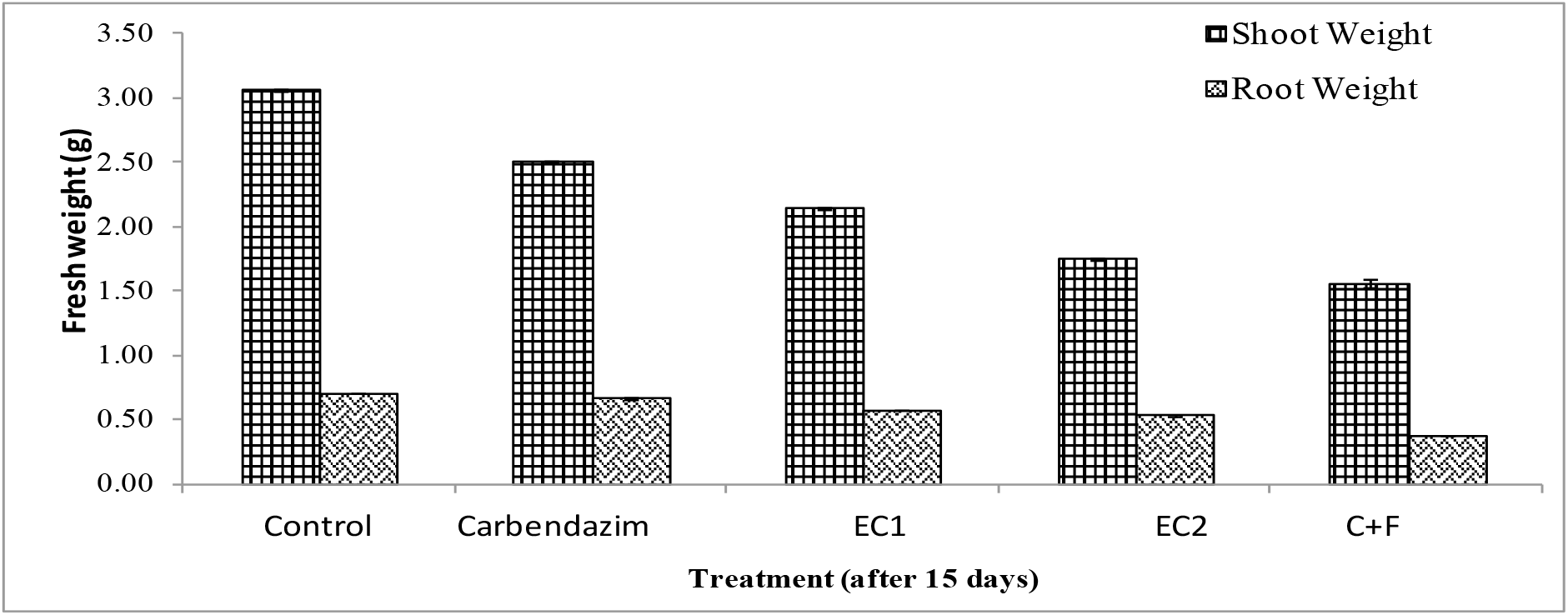
Effect of different doses of CCDM extract on shoot and root fresh weight of tomato plants after 15 days. Control= water; Carbendazim= positive control; EC1=Extract concentration(125μg/mL); EC2= Extract concentration(31.25μg/mL); C+F =negative control (*Fusarium* infested plant without any treatment). Data represents mean ±SE from fivereplicates

**FIGURE 6b.**
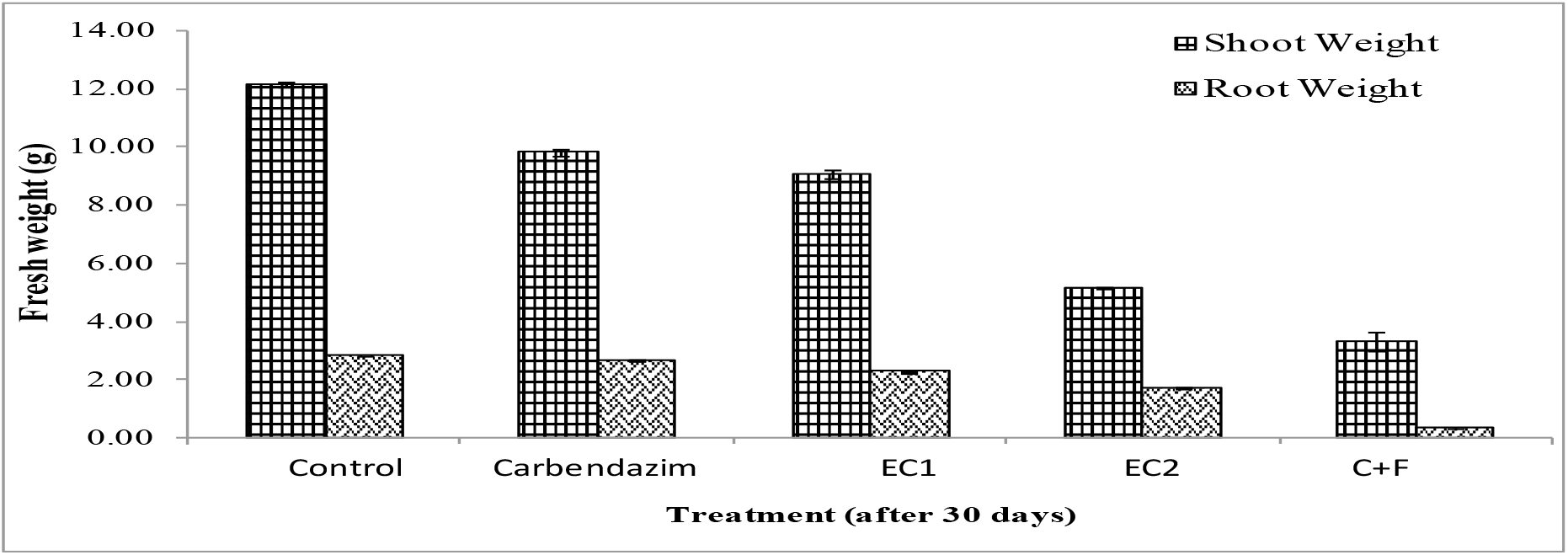
Effect of different doses of CCDM extract on shoot and root fresh weight of tomato plants after 30 days. Control= water; Carbendazim= positive control; EC1=Extract concentration (125μg/mL); EC2=Extract concentration(31.25μg/mL); C+F =negative control (*Fusarium* infested plant without any treatment). Data represents mean ±SE from five replicates

#### 3.4.3 Shoot and root dry weight

Shoot and root dry weight was significantly higher in both EC1 and EC2 treated plants as compared to negative control at all time intervals (Fig 7a, Fig 7b). After 30 days, dry weight of shoot and root was 0.87±0.22g and 0.22±0.005g respectivelyin EC1 treated plants whereas it was 0.94±0.25g and 0.25±0.002g respectively in positive control.

**FIGURE 7a.**
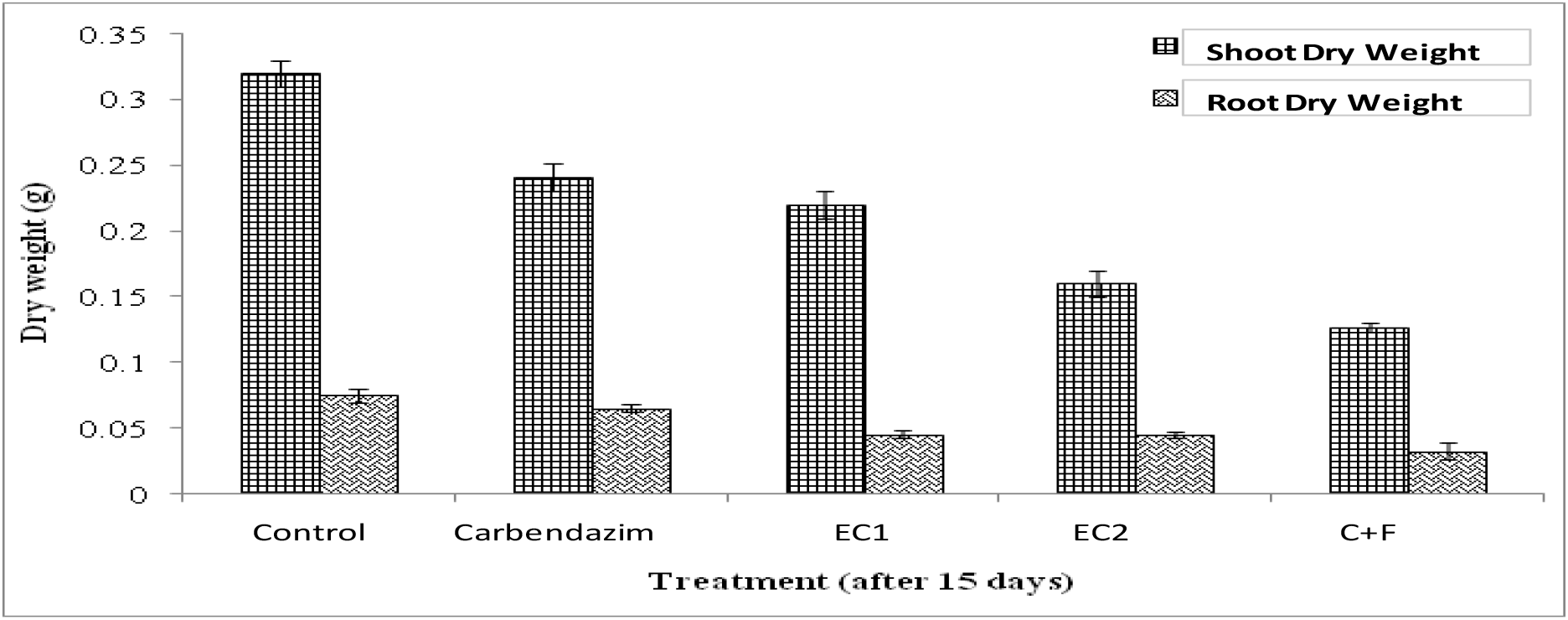
Effect of different doses of CCDM extract on shoot and root dry weight of tomato plants after 15 days. Control=water; Carbendazim= positive control; EC1=Extract concentration(125μg/mL); EC2=Extract concentration (31.25μg/mL); C+F =negative control (*Fusarium* infested plant without any treatment). Data represents mean ±SE from five replicates

**FIGURE 7b.**
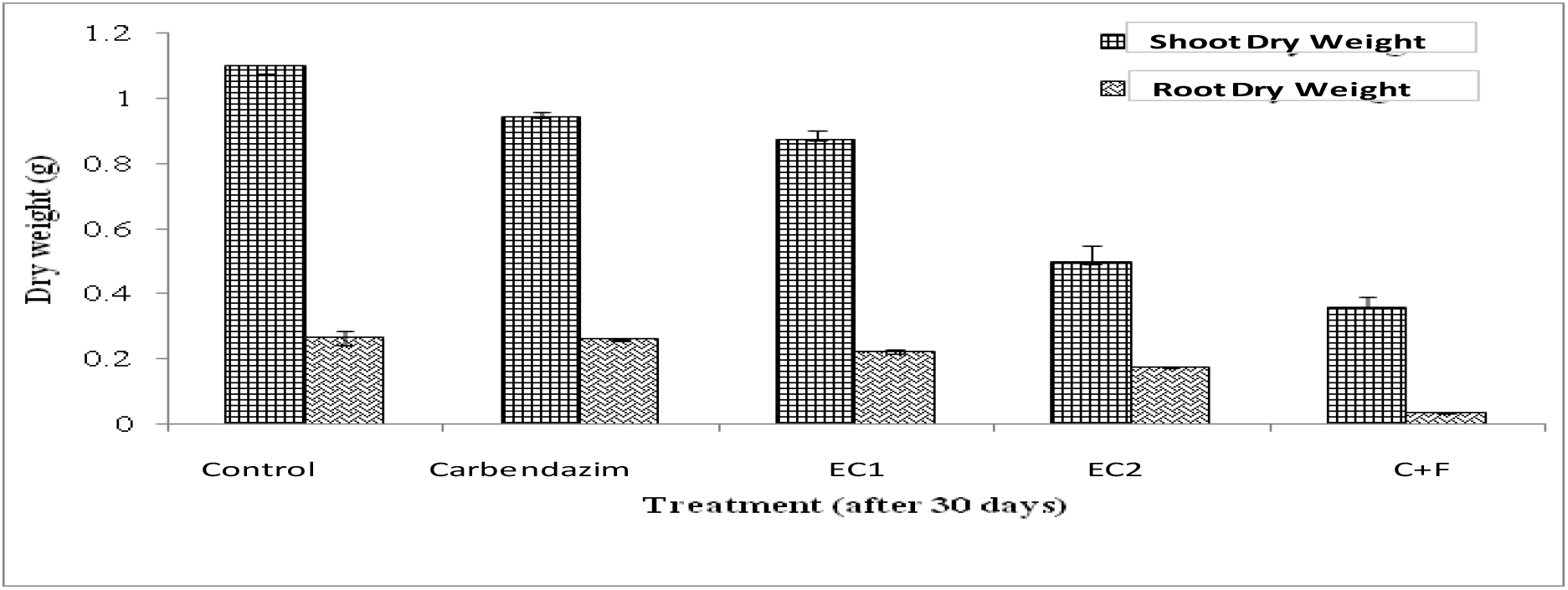
Effect of different doses of CCDM extract on shoot and root dry weight of tomato plants after 30 days. Control=water, Carbendazim= positive control; EC1=Extract concentration (125μg/mL); EC2=Extract concentration(31.25μg/mL); C+F =negative control (*Fusarium* infested plant without any treatment). Data represents mean ±SE from five replicates

### 3.5 Discussion

To the best of our knowledge, no such study is available on the bioefficacyof bryophyte extracts against *Fusarium oxysporum*f. sp.*lycopersici*(FOL) using *in vitro*and *in vivo* approaches. However, reports on *in vitro* antifungal activity of bryophytes against other fungi have been givenby**Mewariet al. (2007), Deora and Guhil(2015**), **Negi et al. (2016a), Negi et al. (2020)**.In the present study, alcoholic extracts of bryophytes showed higher antifungal activity than acetone extracts. Methanol extract of *C. conicum*(CCDM) showed highest mycelial inhibition followed by acetone extract (CCDA) as shown in **Table 1**. Other studies have also shown higher antimicrobial efficacy of alcoholic extract over acetone extract against different microorganisms **(Singh et al. 2011; Kandpal et al. 2016; Negi et al. 2018)**. Variable efficacies of crude organic extracts may be due to the differences in the polarities of the solvents which affect the effective extraction of antifungal compounds in different solvents **(Negi and Chaturvedi, 2016 b)**.

In the present study, significantly higher antifungal activity was exhibited by CCDM extract (plant collected from Dwarahaat, lower altitude) in comparison to CCMM extract (plant collected from Mukteshwar, higher altitude). Similarly, higher antimicrobial activity was also reported in *Dumortierahirsuta* (Sw) Nees collected from lower altitude **(Mukherjee et al. 2012)**.Difference in the bioactivity of the plants growing at different altitudes might be due to variation in accumulation of diverse secondary metabolites influenced by temperature and duration of exposure to UV radiation prevailing at those altitudes **(Zhang and Bjorn 2009)**. Habitat conditions and microniche also govern the responses of different ecotypes occupying diverse habitats.

Since maximum fungal growth inhibition (85.5%) was shown by CCDM extract, hence the same extract was chemically characterized to search for its important chemical constituents responsible for antifungal efficacy against FOL. Chemical characterization of CCDM extract showedthe presence of different bioactive compounds. *viz.*, steroids, fatty acids, sesqueterpenoids, bibenzyls etc. Most of these compounds,like hexadecanoic acid, stigmasteroland δ.5-ergostenolare antifungal in nature **(Ahmed et al. 2010; Mujeeb et al. 2014; Abubakar and Majinda,2016)**. Presence of hexadecanoic acid in organic extract of *C. conicum* was also reported by **Ludwiczuk et al. (2013).**Liverworts also contain lipophilic oil bodies which possessed most characteristic antifungal compounds like bis (bibenzyl) derivatives (riccardin C, riccardin F,isoriccardin C and marchantin A) (**Xie et al. 2010; Asakawa 2013**). Riccardin D isolated from Chinese *Marchantiapolymorpha* showed antifungal activity against fluconazole – resistant *Candida albicans* strains **(Asakawa and Ludwiczuk 2018)**. Riccardin C is one of the most important compounds of liverwortswhich shows strong antifungal activity **(Asakawa et al. 2013)**. Interestingly, GC-MS analysis of CCDM extract in the present study, revealed the presence of very high concentration of riccardin C (64%), which is a major biomarker compound of liverworts having significant antifungal activity. Presence of high concentration of antifungal compounds like riccardin C, fatty acids and steroids in CCDM extract can be directly correlated with its significantly high antifungal activity.Cell membrane damage is the most possible mechanism responsible for antifungal nature of bioactive compounds (**Parvuet al. 2010; Plodpai et al. 2013)**. In the present study, SEM images showedalteration in hyphal morphology which confirmed the fungistatic nature of CCDM extract. Scanning electron microimages clearly showed that EC1concentration of CCDM extract as well as carbendazim treatment (positive control) caused substantial damage to the hyphal structure and hence controlled the growth of FOL mycelia.

*In vitro* antifungal testing of CCDM extract was followed by an *in vivo* glasshouse experiment on tomato plants. The first *in vivo* greenhouse experiment using bryophyte extracts was performed on tomato plants against *Phytophthora infestans* **(Frahm2004)**. In the present study, different effective dosages of *C. conicum* extract were used to control FOLinfection in tomato. Plants grown in EC1 amended pots attained significantly higher shoot and root height, shoot and root fresh weight and dry weight as compared to EC2 and the negative control. All these parameters were quite close to the positive control and indicatedsignificant biocontrol potential of CCDM. Fusarium wilt caused blockage inside the xylem vessels by mycelia producing microconidia, which travel upward in the transpiration stream **(Okungbowa and Shittu2012)**. Interruption of the xylem vessels and transpiration stream caused wilting in fusarium infested plants. It also caused poor development of lateral roots because of high infection rate **(Loganathan et al. 2009)**.Poor rootand shootgrowth was also seen in fusarium infested plants in the present study. However, tomato treated plants with EC1 and EC2 dosage of CCDM could overcome the biological stress caused by the fungus and showed significantly higher root and shoot growth parameters compared to fusarium control(C+F). Here, in the study, EC1 concentration of CCDM extract was found to be more effective than EC2. Chemical characterization of CCDM extract revealed presence of large number of antifungal compounds. One of the major bioactive compoundof CCDM extract was riccardin C which is a well known antifungal compound and contributes significantly to the antifungal potential of *C. conicum*.

### 3.6 Conclusion

The present study was aimed to find an ecofriendly biological control of *Fusarium oxysporum*f. sp. *lycopersici* (FOL) from lesser known cryptogamic plants *viz*., bryophytes. The first step to meet the objective was finding a potent bryophyte *viz*., *Conocephalumconicum*, which inhibited the mycelia growth of FOL (85 % per cent inhibition) in a laboratory test. The second step was to find out the effective dosage of the biocontrolling botanical. The effective concentration (125μg/ml) of the methanolic extract of *C. conicum* disrupted the hyphalstructure and emerged as the most potent dosage to control fusarium wilt under glasshouse conditions as well. To the best of our knowledge, this is the first study to report theuse of organic extracts of *C. conicum* to control *F. oxysporum*f. sp. *lycopersici*using*in vitro* and *in vivo* approaches. Present studyproved the antifungal potential of crude methanol extract of *C. conicum*(owing to the presenceof riccardin c and other antimicrobial compounds). Based on the present study, *C. conicum* can be utilized as an environment friendly botanical fungicide providing a promising alternative to chemicals. However, still extensive field and advance ultra structuralstudies are needed to understand the molecular mode of interaction between the bioactive compound and the pathogen.Procuring enough quantity of the material is another challenge that can be tackled by scaling up their propagation in bioreactors.

## Acknowledgements

We are thankful to Dr. S.D. Tewari, MMV College Haldwani, Kumaon University, Nainital for identification of plants. The generous help in authentication of plant identity rendered by Dr. A.K. Asthana and Dr. Vinay Shahu, NBRI, Lucknow is also duly acknowledged. Authors are highly thankful to Uttarakhand Council of Science and Technology for funding this work. The work was supported by research grant (File No. UCS&T/R&D/LS-29/11-12/4345/2 dated 14-(03-2012) offered by Uttarakhand Council of Science & Technology to PC. KN was awarded research fellowship by the council.

